# Working with Zika and Usutu Viruses in Vitro

**DOI:** 10.1101/040139

**Authors:** Kelli L. Barr, Benjamin D. Anderson, Maureen T. Long

**Affiliations:** Department of Infectious Diseases & Pathology, College of Veterinary Medicine, University of Florida, 2015 SW 16th Av2015 SW 16th Ave, Gainesville, FL, USA.; Division of Infectious Disease, School of Medicine and Global Health Institute, Duke University, Durham, North Carolina, USA

**Keywords:** Usutu virus, Zika virus, flavivirus, host range, fusion, cytopathic effects

## Abstract

Usutu and Zika viruses are emerging arboviruses of significant medical and veterinary importance. These viruses have not been studied as well as other medically important arboviruses such as West Nile, dengue, or chikungunya viruses. As such, information regarding the behavior of Zika and Usutu viruses in the laboratory is dated. Usutu virus re-emerged in Austria in 2001 and has since spread throughout the European and Asian continents causing significant mortality among birds. Zika virus has recently appeared in the Americas and has exhibited unique characteristics of pathogenesis, including birth defects, and transmission. Information about the characteristics of Usutu and Zika viruses are needed to better understand the transmission, dispersal, and adaptation of these viruses in new environments. Since their initial characterization in the middle of last century, technologies and reagents have been developed that could enhance our abilities to study these pathogens. Currently, standard laboratory methods for these viruses are limited to 2-3 cell lines and many assays take several days to generate meaningful data. The goal of this study was to characterize these viruses in cell culture to provide some basic parameters to further their study. Cell lines from 17 species were permissive to both Zika and Usutu viruses. These viruses were able to replicate to significant titers in most of the cell lines tested. Moreover, cytopathic effects were observed in 8 of the cell lines tested. The data show that, unlike other flaviviruses, neither Zika nor Usutu viruses require an acidic environment to fuse with a host cell. This may provide a tool to help characterize events or components in the flavivirus fusion process. These data indicate that a variety of cell lines can be used to study Zika and Usutu viruses and may provide an updated foundation for the study of host-pathogen interactions, model development, and the development of therapeutics.

**Author Summary:** Usutu and Zika viruses are arboviruses of identified in significant medical and veterinary outbreaks in recent years. Currently, standard laboratory methods for these viruses are limited to 2-3 cell lines and basic viral characterization has not been performed since the mid-20^th^ century. Zika and Usutu viruses were characterized in cell culture. The data show that a variety of cell lines can be used to study the viruses. Neither Zika nor Usutu viruses require an acidic environment for host cell infection.

## Introduction

Usutu virus (USUV), first identified in South Africa in 1959, is a flavivirus belonging to the Japanese encephalitis complex [1,2]. In 2001, USUV emerged in Austria and spread throughout the European and Asian continents [3-10]. Unlike USUV circulating in Africa, the new emergent strains caused significant mortality among Old World blackbirds, owls, and other wild and captive birds [3,11].

The host range of USUV includes primarily Culex mosquitoes, birds, and humans [1] and is most often transmitted between avian reservoir hosts and mosquitoes in a sylvatic transmission cycle. Infections with USUV are usually non-pathogenic in humans. Other than birds, evidence for USUV infection has been found in humans and horses [12-14]. Several human cases have been identified in Europe and Croatia [15-17]. Recently, USUS has been linked to neuroinvasive infections in 3 patents from Croatia [10] and has been detected in horses in Tunisia [14].

Zika virus (ZIKA) is an emerging, medically important arbovirus. It is classified as a flavivirus and is descendent from Yellow fever virus [18]. Like many other tropical arboviruses, human infection with ZIKA typically presents as an acute febrile illness with fever, rash, headache, and myalgia. The flavivirus, Dengue virus (DENV) and the alphavirus, chikungunya virus (CHIK) produce similar symptoms to ZIKA but are more commonly diagnosed. The high seroprevelance of ZIKA antibodies in human populations in Africa and Asia suggests the misdiagnosis of ZIKA for other arboviral illnesses is an ongoing problem [19]. There are two geographically distinct lineages of circulating ZIKA; African and Asian [19]. The Asian lineage has recently emerged in Micronesia where it was the cause of a large outbreak in 2007 [20] and currently in the Americas [21].

The natural hosts of ZIKA include humans, primates, and *Aedes* mosquitos [22-25]. Though no solid evidence exists of non-primate reservoirs of ZIKA [26], antibodies to ZIKA have been detected in elephants, goats, lions, sheep, zebra, wildebeests, hippopotamuses, rodents, and other African ruminants [27,28].

There are several characteristics of ZIKA that distinguish it from other medically important arboviruses. In recent outbreaks in French Polynesia, ZIKA exhibited increased pathogenicity and atypical symptoms including respiratory involvement and conjunctivitis [20,29]. A ZIKA strain acquired in Senegal during 2008 exhibited the ability to spread from human to human through sexual transmission [30]. Zika virus has been detected in cell nuclei, unlike other flaviviruses that are confined to the cellular cytoplasm [31]. During the current outbreak in the Americas, ZIKA has been linked to serious medical conditions. Maternal-fetal transmission of ZIKA has resulted in microcephaly and other brain abnormalities [32, 33]. In five of 49 cases, ZIKA was detected in the brain of the children [33]. Guillain-Barré (GB) syndrome is also being associated with these outbreak isolates [34], however GB is one of the most common differential diagnoses for West Nile virus.

Due to their lack of apparent clinical importance in humans and animals, USUV and ZIKA have not been studied to the same degree as other, wide spread flaviviruses such as West Nile virus, DENV or CHIK. While research in serology and genetic characterization are underway [19,20,34], the recent changes in biology and distribution of these viruses warrant further investigation as many questions regarding the basic biology and ecology of ZIKA and USUV remain unanswered. To better understand the characteristics of USUV and ZIKA *in vitro*, we investigated the permissiveness of several cell lines and determined the basic fusion requirements of these viruses in cell culture.

## Materials and Methods

### Cells and viruses

Seventeen cell lines were obtained from the ATCC (Manassas, VA) and included TB 1 Lu, DF-1, Sf 1 Ep, EA.hy.926, CRFK, E.Derm, FoLu, Pl 1 Ut, OHH1.K, OK, DN1.Tr, PK(15), LLC-MK2, BT, MDCK, WCH-17, Mv1 Lu (Table 1). These lines were selected to include representatives of species found only in the Americas; specifically, North America. All cell lines were cultured in Dulbecco's modified eagle medium (DMEM) supplemented with 10% (v/v) fetal bovine serum (FBS), 1% (v/v) L-glutamine, 1% (v/v) non-essential amino acids (NEAA), 1% (v/v) sodium pyruvate, 100 U/ml penicillin, 100 μg/ml streptomycin, and housed in a 37°C incubator with 5% CO2.

**Table 1.**
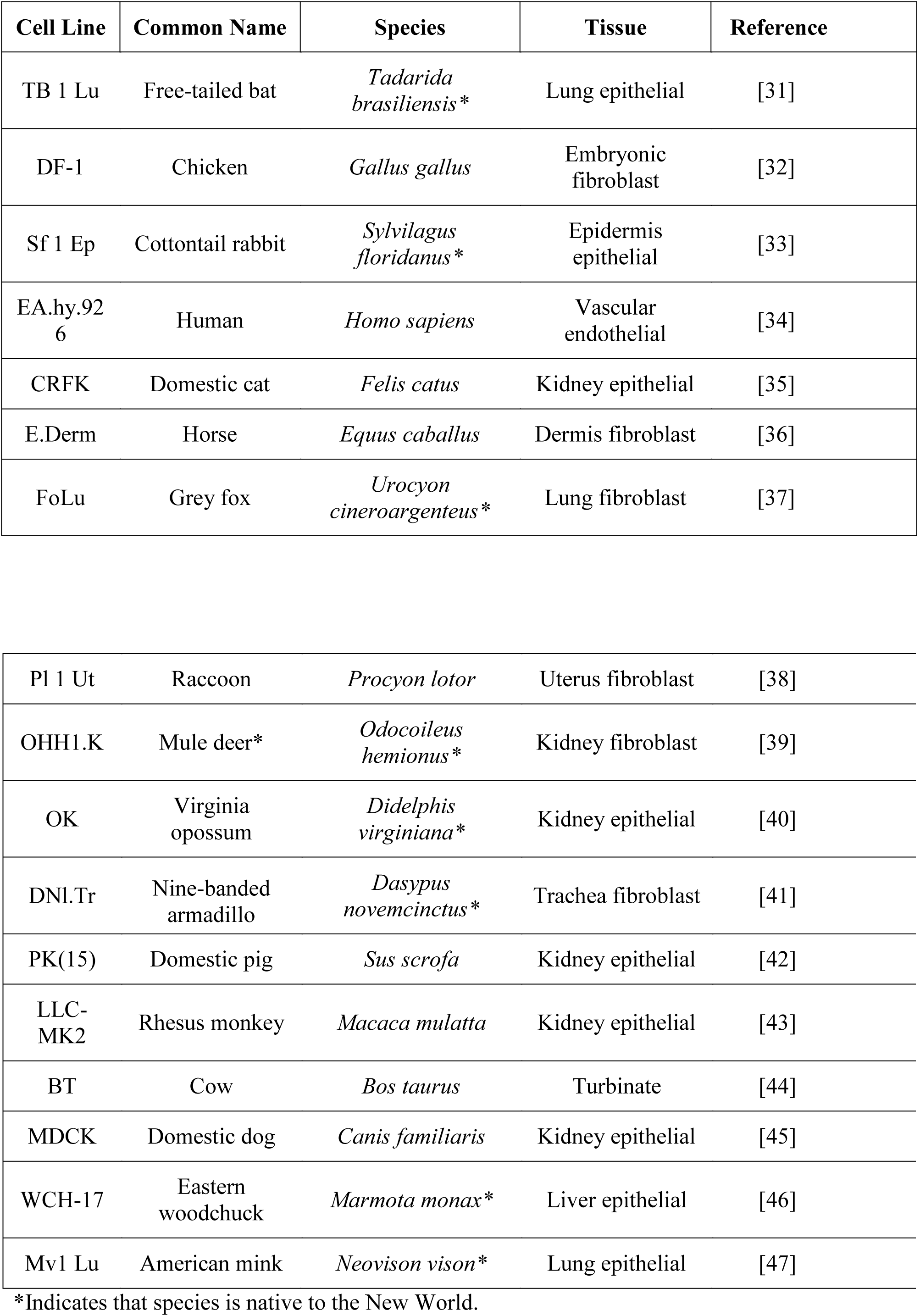
Cell lines used for characterization of Usutu and Zika viruses.

| Cell Line | Common Name | Species | Tissue | Reference |
| --- | --- | --- | --- | --- |
| TB 1 Lu | Free-tailed bat | *Tadarida brasiliensis* * | Lung epithelial | [31] |
| DF-1 | Chicken | *Gallus gallus* | Embryonic fibroblast | [32] |
| Sf 1 Ep | Cottontail rabbit | *Sylvilagus floridanus* * | Epidermis epithelial | [33] |
| EA.hy.926 | Human | *Homo sapiens* | Vascular endothelial | [34] |
| CRFK | Domestic cat | *Felis catus* | Kidney epithelial | [35] |
| E.Derm | Horse | *Equus caballus* | Dermis fibroblast | [36] |
| FoLu | Grey fox | *Urocyon cinereoargenteus* * | Lung fibroblast | [37] |
| Pl 1 Ut | Raccoon | *Procyon lotor* | Uterus fibroblast | [38] |
| OHH1.K | Mule deer* | *Odocoileus hemionus* * | Kidney fibroblast | [39] |
| OK | Virginia opossum | *Didelphis virginiana* * | Kidney epithelial | [40] |
| DNI.Tr | Nine-banded armadillo | *Dasypus novemcinctus* * | Trachea fibroblast | [41] |
| PK(15) | Domestic pig | *Sus scrofa* | Kidney epithelial | [42] |
| LLC-MK2 | Rhesus monkey | *Macaca mulatta* | Kidney epithelial | [43] |
| BT | Cow | *Bos taurus* | Turbinate | [44] |
| MDCK | Domestic dog | *Canis familiaris* | Kidney epithelial | [45] |
| WCH-17 | Eastern woodchuck | *Marmota monax* * | Liver epithelial | [46] |
| Mv1 Lu | American mink | *Neovison vison* * | Lung epithelial | [47] |

\*Indicates that species is native to the New World.

Usutu virus (SAAR-1776), ZIKA (MR766), Yellow Fever virus (17D), Sindbis virus (EgAr 339), CHIK (181/25), DENV-1 (H87), DENV-2 (NGC), DENV-3 (HI), and DENV-4 (H241) were obtained from the World Reference Center for Emerging Viruses and Arboviruses (Robert Tesh, UTMB, Galveston, TX). West Nile virus (NY99) was obtained from the University of Florida (Maureen Long).

### Infection of cells with viruses

All infections were performed using 12 or 24-well standard cell culture plates seeded with cells which had reached a 90% confluence upon infection. Individual wells were inoculated with 1,000 infectious units (IU) of virus in MEM and then rocked at 37°C for one hour after which the inoculum was removed, rinsed twice with sterile PBS, overlaid with 1 ml of DMEM (10% FBS, 1% glutamine, 1% NEAA, 100mg/ml penicillin/streptomycin, 1% sodium pyruvate) and incubated at 37°C incubator with 5% CO_2_. Culture supernatants were collected at 1 and 72 hours post-inoculation (PI).

### Visualization of cytopathic effects

Cells were examined daily for cytopathic effects (CPE). All cell lines were allowed to develop CPE for 7 days post infection. Cells were stained using 70% ethanol containing 1% wt/vol crystal violet. Plates were incubated for 15 minutes at 22°C after which the fixative was decanted. The plates were rinsed in cold tap water and dried overnight at room temperature. Images were obtained using Micron imaging software (Westover Scientific) and an inverted microscope at 40X magnification.

### Primer design for qRT-PCR

Primers for USUV were designed against the USU181 sequence (Genbank accession: JN257984) and amplify a 104 base pair fragment of the envelope protein gene starting at nucleotide position 239 and ending at position 342. Primers for ZIKA were designed against the MR766 strain (Genbank accession: AY632535) and amplify a 128 base pair fragment of the envelope glycoprotein starting at nucleotide position 1398 and ending at position 1525. Blasts for these primer sequences showed sequence homology to multiple strains of the reference virus but no homology to other viruses. The USUV primer set could detect as few as 10 IU per mL and the ZIKA primer set was able to detect as few as 100 IU/mL. Both primer sets did not amplify other arboviruses tested including: West Nile virus, Sindbis virus, Yellow fever virus, DENV serotypes 1-4, and CHIK. Sequences for the primer sets are listed below:

Usutu_Forward (5’-AGCTCTGACACTCACGGCAACTAT-3’)

Usutu_Reverse (5’-TCACCCATCTTCACAGTGATGGCT-3’)

Zika_Forward (5’-TATCAGTGCATGGCTCCCAGCATA-3’)

Zika_Reverse (5’-TCCTAAGCTTCCAAAGCCTCCCAA-3’)

### Virus detection via real-time RT-PCR

Viral RNA was extracted from cell culture supernatant using the Ambion MagMax-96 extraction kit (Life Technologies: Grand Island, NY) per manufacturer’s instructions. Quantitative, real time, reverse transcriptase polymerase chain reaction (qRT-PCR) was conducted with BioRad Superscript One Step SYBR Green qRT-PCR kit (Winooski, VT). The following cycling conditions were employed: reverse transcription at 50°C for 10 min, denaturation at 95°C for 5 min, followed by 40 cycles of denaturation and amplification at 95°C for 10 sec and 55°C for 30 sec. Cycle threshold (Ct) values were used to estimate relative viral titers of infected cell lines according to a standard curve created using a serial dilution of known viral concentrations. Results are expressed as the average of 3 independent trials amplified in duplicate.

A series of controls were performed for each cell line in order to identify true positives not related to background. A no-template control and a no-primer control were performed to verify that the reagents and equipment were working as expected. A positive virus control was used to verify that the PCR primers were functioning as expected. A non-infected cell culture supernatant control was included to verify that there was no increase in non-specific binding from the PCR primers that could cause a higher background signal. Finally, the cell culture supernatant collected 1 hour PI to ensure that qRT-PCR results, 72 hours PI, were not convoluted by input virus.

### Fusion inhibition assay

To determine if virus infectivity was pH dependent, the pH drop that occurs in the cellular endosome during viral fusion was inhibited as previously described [52]. Briefly, LLC-MK2 cells were pre-treated with blocking media (DMEM, 0.2% BSA, 10 mM Hepes, 50 mM NH4Cl pH8) for two hours at 37°C, and the cells were then infected with 10,000 IU virus in the presence of 50 mM NH4Cl and incubated for 1 hour at 37°C. The cultures were then rinsed with PBS and incubated for an additional two hours at 37 °C in blocking media after which the media was replaced with DMEM with 10% FBS. Cell culture supernatants were harvested 48 hours PI to determine extracellular virus yields by qRT-PCR. RNA extractions and qRT-PCR were performed on the cell monolayers. Results are expressed as the average of three independent trials amplified in duplicate.

### Virus binding assay

To determine if cell resistance to USUV or ZIKA was binding dependent, a virus: cell binding assay was performed as previously described [52] with RNA extractions and qRT-PCR performed on the cell monolayers. Results are expressed as the average of 3 independent trials amplified in duplicate.

## Results

### USUV and ZIKA replicate in multiple cell lines

Of the 17 cell lines tested for USUV infection, 16 showed quantifiable Ct values based upon qRT-PCR data at 72 hours PI (Figure 1). All cell lines except WHC-17 (*Marmota monax*) produced at least 10^3^ relative infectious units. The cell lines BT (*Bos Taurus)*, PK(15) (*Sus scrofa*), FoLu (*Urocyon cineroargenteus*), CRFK (*Felis catus*), OHH1.K (Odocoileus hemionus hemionus), DF-1 (*Gallus gallus*), MDCK (*Canis familiaris*), and OK (*Didelphis marsupialis virginiana*) were able to replicate USUV as well as or better than the LLC-MK2 cell line (Figure 1). The OK cells were able to produce over 10^7^ relative infectious units; over one and a half logs more USUV than LLC-MK2 cells (Figure 1), indicating their potential use for virus culture.

**Figure 1.**
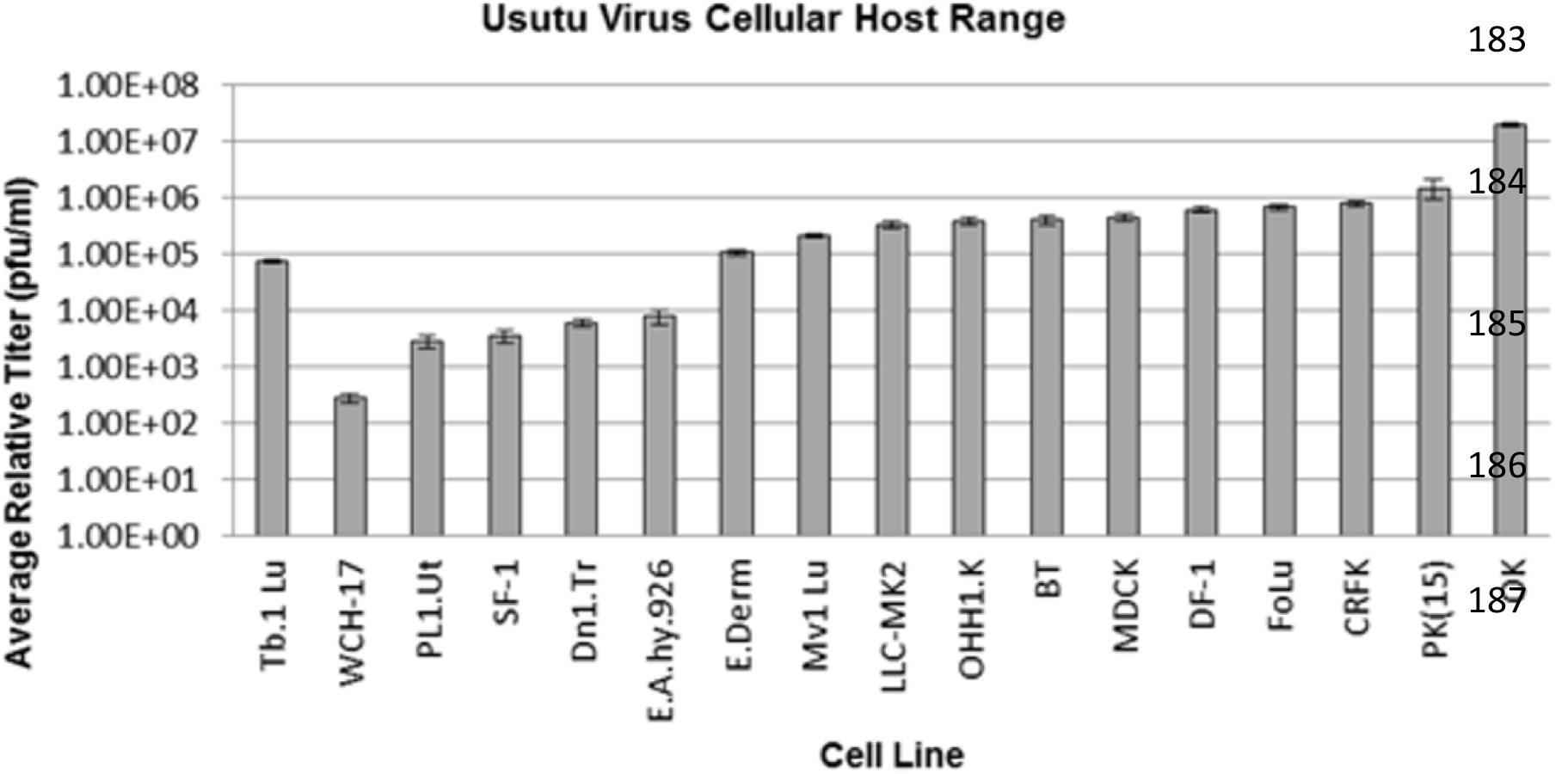
The host range of USUV in cell culture. Mean relative titers of USUV ± SEM produced from cell culture supernatants from 17 cell lines collected at 72 hours post-infection. Relative viral titers of infected cell lines were calculated according to a standard curve created using a serial dilution technique of known viral concentrations.

Of the 17 cell lines tested for ZIKA infection, 15 showed quantifiable Ct values based upon qRT-PCR data at 72 hours PI (Figure 2). All cell lines except WHC-17 (*Marmota monax*) and TB 1 Lu (*Tadarida brasiliensis*) produced at least 10^4^ relative infectious units. The cell lines E.Derm (*Equus caballus*), PK(15), FoLu, CRFK, and OK were able to replicate ZIKA as well as or better than the LLC-MK2 cell line (Figure 2). The OK cells were able to produce over 10^6^ relative infectious units; up to a log more ZIKA than LLC-MK2 cells (Figure 2), indicating their potential use for virus culture.

**Figure 2.**
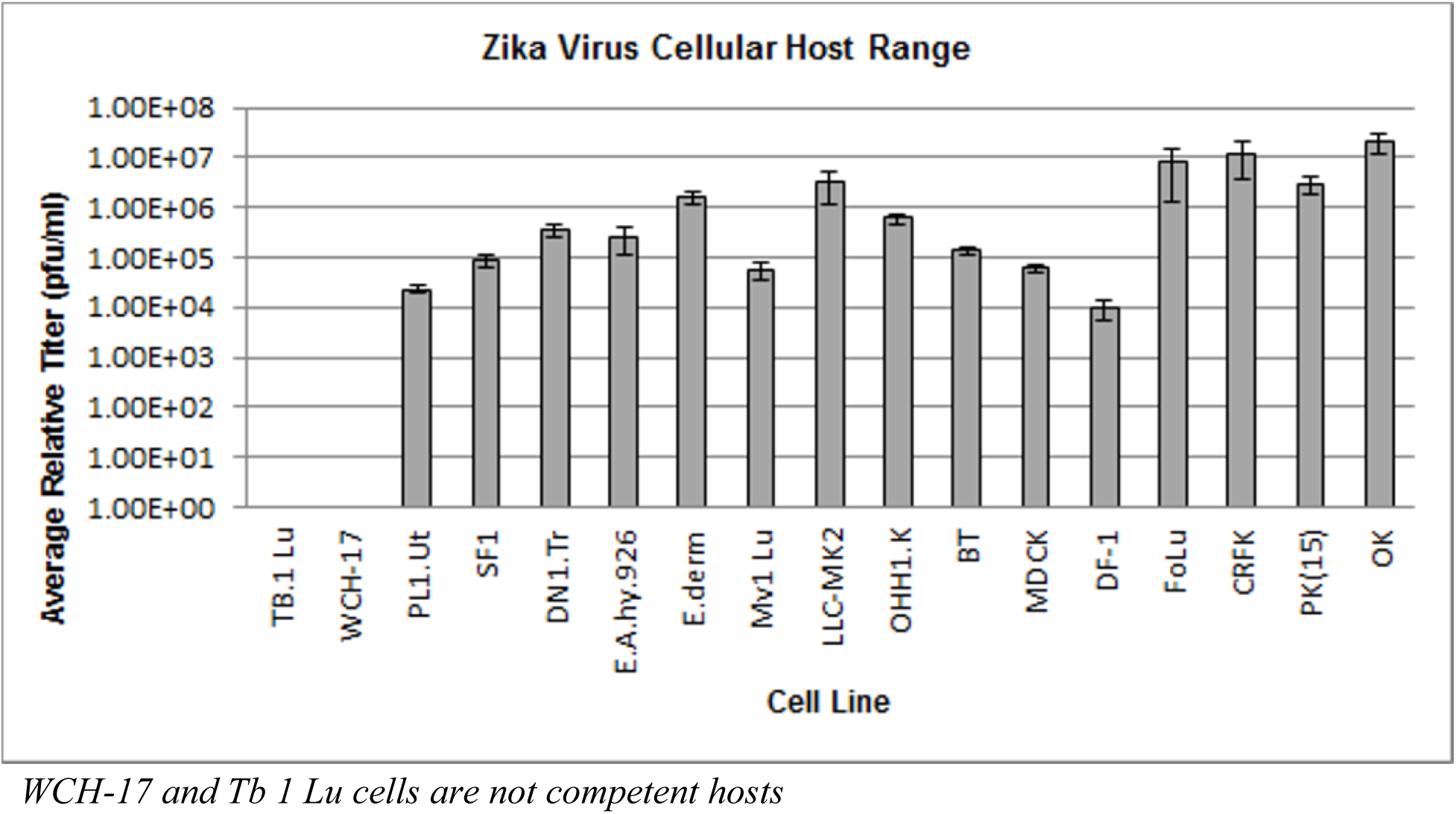
The host range of ZIKA in cell culture. Mean relative titers of ZIKA ± SEM produced from cell culture supernatants from 17 cell lines collected at 72 hours post-infection. Relative viral titers of infected cell lines were calculated according to a standard curve created using a serial dilution technique of known viral concentrations.

### WCH-17 and Tb 1 Lu cells are not competent hosts

Usutu was not detected in low quantities in WCH-17 (*Marmota monax*) cells and ZIKA was not detected in Tb 1 Lu (*Tadarida brasiliensis*) or WCH-17 cells via qRT-PCR nor was CPE evident. A virus: cell binding assay was performed in order to determine if cell receptors were present that would allow ZIKA or USUV to attach to the Tb 1 Lu or WCH-17 cell surface. The Ct values for all treatments express the amount of virus present in the sample. The statistical similarity of the data suggests that both ZIKA and USUV bind to WCH-17 cells and ZIKA binds to Tb 1 Lu cells as efficiently as they bind to the LLC-MK2 control cells (Table 2).

**Table 2.** Ct values as determined by qRT-PCR of ZIKA and USUV after binding to LLC-MK2, Tb 1 Lu, and WCH-17 cells. The lack of significant difference between Ct values of the three cell lines indicate that both Zika and Usutu viruses bind to WHC-17 and/or Tb1. Lu cells as efficiently as they bind to the LLC-MK2 control cells.

|  | LLC-MK2 | WCH-17 | Tb1. Lu |
| --- | --- | --- | --- |
| **ZIKA** | 20.09 ($\pm 0.095$) | 20.19 ($\pm 0.26$) | 20.57 ($\pm 0.08$) |
| **USUV** | 20.95 ($\pm 0.095$) | 20.99 ($\pm 0.095$) | NA |

### USUV and ZIKA produce cytopathic effects in multiple cell lines

Cytopathic effects were observed in CRFK, Dn1.Tr (*Dasypus novemcinctus*), Sf 1 Ep (*Sylvilagus floridanus*), PK(15), FoLu, Mv 1 Lu (*Neovison vison*), OHH1.K, and OK cell lines from both ZIKA and USUV infection. Forms of CPE caused by USUV included the formation of koilocytes (Figure 3a, 3b, 3c), cellular enlargement (Figure 3d, 3f), rounding (Figure 3f, 3h), focal degeneration (Figure 3g), and pyknosis (Figure 3d, 3e, 3h). Forms of CPE caused by ZIKA included the formation of koilocytes (Figure 3b, 3f, 3h), cellular enlargement (Figure 3b, 3h), focal degeneration (Figure 3c, 3d, 3f, 3g), and pyknosis (Figure 3a, 3d, 3e).

**Figure 3.**
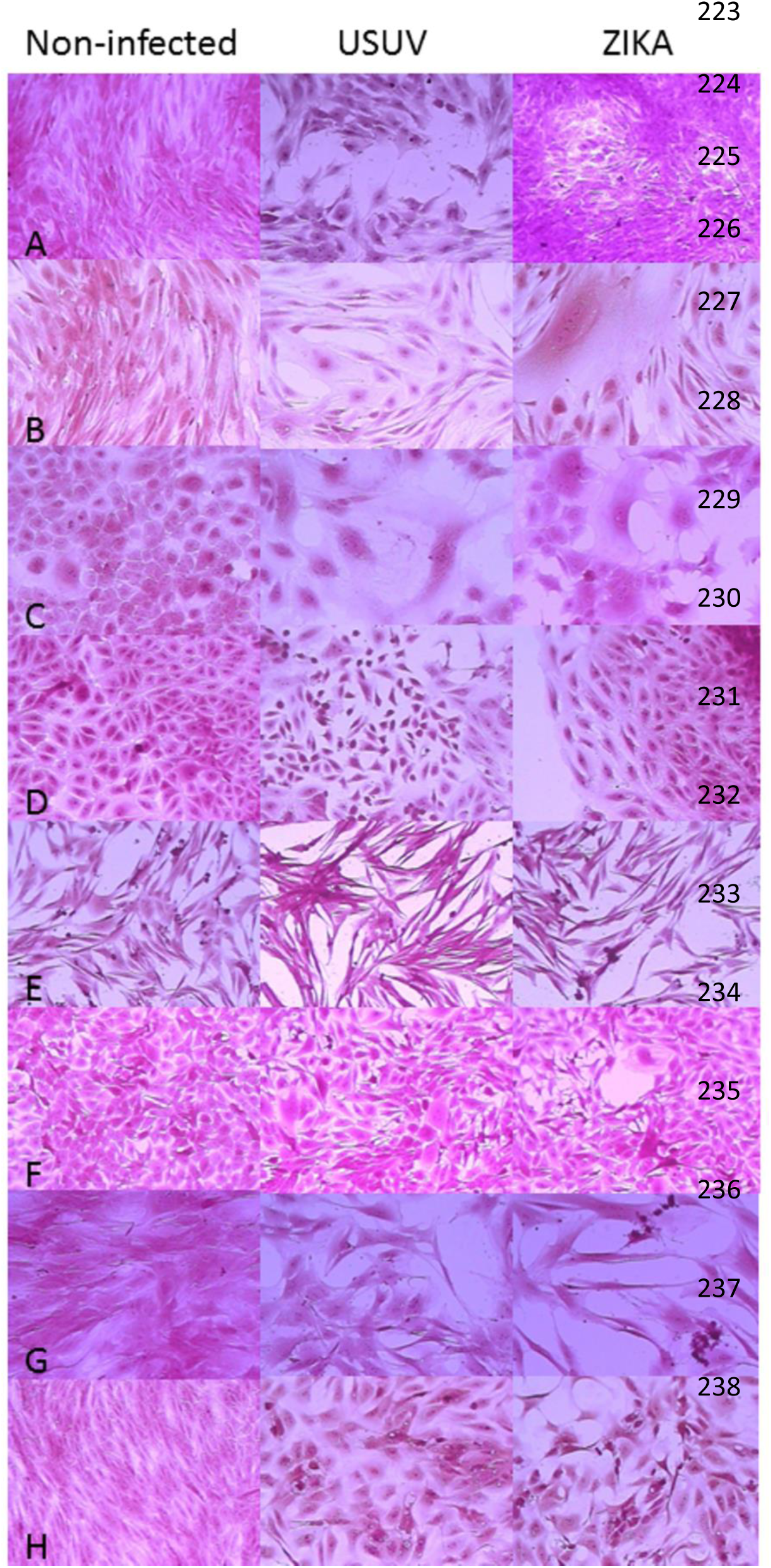
The cytopathic effects of USUV and ZIKA. Cytopathic effects of Zika and Usutu viruses were visualized a 40X magnification on an inverted microscope. Cytopathic effects were observed for both viruses in (a) CRFK, (b) Dn1.Tr, (c) Sf 1 Ep, (d) PK(15), (e) FoLu (f) Mv 1 Lu, (g) OHH1.K, and (h) OK cells.

### FoLu, USUV and ZIKA do not require pH mediated fusion

The ability of USUV and ZIKA to fuse with a host cell was evaluated by blocking the drop in pH that occurs in the cellular endosome which has been shown to induce conformational changes in the viral envelope necessary for fusion. The performance of DENV-2 was evaluated in tandem as a control as it has been shown that DENV, and other flaviviruses, require an acidic pH to fuse with a host cell [52, 54]. The data show that USUV and ZIKA were able to fuse with the target cells in the absence of acidic pH as well as the control (acidic) treatment (Table 3). The Ct values for USUV and ZIKA were statistically similar despite pH level. The Ct values for DENV (the control virus) were significantly lower for cells treated with and acidic pH. This indicates that more virus was present in the cells than what was detected in the cells infected with DENV in an environment of basic pH.

**Table 3.** Ct values as determined by qRT-PCR of USUV, ZIKA, and DENV fusion with LLC-MK2 cells in the presence and absence of and acidic endosome. Usutu and Zika viruses did not exhibit inhibited host cell fusion in the presences of a basic pH whereas the fusion of dengue virus, the control, was significantly inhibited in the presence of a basic pH.

|  | Acidic pH | Basic pH |
| --- | --- | --- |
| USUV | 13.34 ($\pm 1.11$) | 14.58 ($\pm 0.44$) |
| ZIKA | 7.8 ($\pm 0.2$) | 8.87 ($\pm 0.54$) |
| DENV | 17.32 ($\pm 0.3$) | 22.18 ($\pm 0.15$) |

## Discussion

Though evidence of ZIKA infection has been found in non-primate species, the host range for ZIKA, both *in vitro* and *in vivo*, has not yet been explored. Preliminary studies using the 2001 USUV emergent strain indicated that the virus could infect several species in cell culture [55]. For this experiment, we sought to examine the host range of the prototype ZIKA and USUV isolates in cell culture. Eighteen distinct cell lines were selected from the inventory at the American Type Culture Collection (ATCC) (Manassas, VA) and included species that were only found in the Americas. Cell lines were selected based on the susceptibility of the host species to flaviviral infection and utility of the cell line in virus research. ZIKA and USUV are Old World viruses and as such have not encountered New World species like opossum, armadillo, North American mule deer, raccoon, gray fox, and cotton-tail rabbit. Most of these animals are peridomestic and inhabit the same environment as the mosquito vectors. It is of particular interest that both USUV and ZIKA replicate well in cells from many domestic and peridomestic animals. These animals may be susceptible to clinical disease and if viremia is high enough, they may serve as reservoirs or hosts. Viral transmission and encroachment may reflect that of the West Nile introduction to the United States in 1999. The data agree with other work that shows USUV can infect PK (15), MDCK, and primate cells [55] and suggest that the USUV prototype strain may behave similarly in cell culture to the emergent strains of the virus. In addition, This work agrees with recent work showing that DENV can replicate in a variety of cell lines [56, 57]

The data show that ZIKA and USUV bind to WCH-17 cells and ZIKA binds to Tb 1 Lu cells as efficiently as the LLC-MK2 control cells. This suggests that USUV or ZIKA infection of WCH-17 and Tb 1 Lu cells may be inhibited during the virus: cell fusion or viral replication process.

Notably, WCH-17 cells are infected with hepatitis B virus which may be a contributing factor to the inability of these viruses to establish an infection in this cell line.

In addition to replicating in various cell lines, USUV induced cytopathic effects (CPE) in 8 of the 16 positive cell lines. The characteristics of CPE caused by a flavivirus vary in accordance to the host cell [58] and are dependent on various factors including host genetics, viral receptors, immune-response, and defective virus particles [58]. Previous studies on the 2001 USUV emergent strain indicated that CPE was induced in PK (15), Vero, and GEF (goose embryo fibroblast) cells [55]. The range and extent of CPE observed suggests that these cell lines may be useful for virus culture and viral titer studies such as TCID50 and plaque reduction neutralization tests.

The entry of a flavivirus into a host cell is dependent upon clathrin-mediated endocytosis [59-62]. Fusion of the viral membrane with the host cell requires conformational changes to the viral envelope glycoprotein that are induced by a low-pH [63-66]. Though, alternative infectious pathways for flaviviruses have been described [reviewed by Smit et al. [52,67], it is agreed that an acidic pH is necessary for successful flaviviral fusion with the host cell [45-50]. The ability of a flavivirus to fuse with a target cell is a function of the tertiary protein structure of the envelope glycoprotein. The conformation of the glycoprotein is based on the nucleic acid sequence of the glycoprotein gene. It has been shown that differences in the amino acid sequences of the envelope glycoprotein are associated with significant changes in pathogenicity, clinical presentation, resistance, and hydrophobic/hydrophilic properties [68-71]. The data for DENV agree with other research as its ability to infect a target cell was significantly inhibited in the absence of an acidic environment within the endosome [52]. It may be that unique features are present on the envelope glycoprotein of USUV and ZIKA that are involved in the fusion process that are permitting these viruses to fuse in a non-acidic environment.

Research has shown that USUV is genetically distinct from other flaviviruses [72]. Different strains of USUV have been shown to differ by as much as 5% in amino acid sequence [8]. These amino acid substitutions may influence virulence and other characteristics of USUV [8,72,73]. ZIKA too, is genetically distinct from other flaviviruses, and different strains of ZIKA have been shown to differ by as much as 11.7% in nucleotide sequence [19]. Moreover, significant amino acid deletions have been identified at glycosylation sites of the envelope glycoprotein in some strains of ZIKA [19], which may influence virulence or other characteristics of the virus [74].

USUV and ZIKA may achieve their broad host range by exploiting alternative infectious entry pathways. Cellular membrane components such as clathrin, dynamin, actin, and lipids have been shown to be involved with viral entry into the host cell cytoplasm [52,75-79]. The impact of these various components on virus entry is has been shown to be host specific for DENV [52] and may be contributing to the ability of USUV or ZIKA to establish infection in a wide variety of cell lines.

## Conclusions

The data herein indicate that several cell lines can be used to culture and study USUV and ZIKA and that an acidic environment is not required in the cellular endosome to achieve successful fusion with a host cell. The susceptibility for certain cell lines to USUV and ZIKA may provide a tool for characterizing these viruses and may provide an *in vitro* platform for the study of host-pathogen interactions, model development, and the development of therapeutics. The unique fusion requirements of USUV and ZIKA may be useful in understanding flaviviral infection and may identify novel targets for the development of interventions. Though these experiments raised some provocative questions, there were some limitations to this study, which should be addressed. For instance, the broad host infectivity observed may be a function of the virus strains that were used for the experiments. These strains may not accurately reflect the characteristics of USUV or ZIKA currently circulating, or that of other laboratory-adapted strains. Finally, the behavior of USUV and ZIKA in the laboratory does not reflect the behavior of these viruses in their natural environment.

## Acknowledgments

We would like to thank Dr. Robert Tesh at WRCEVA for sharing viruses. We thank Dr. Gregory C. Gray for the generous use of his RT-PCR equipment and BSL2 facilities.

## References

1. Nikolay B, Diallo M, Boye CS, Sall AA. Usutu virus in Africa. Vector Borne Zoonotic Dis. 2011;11(11):1417–23. doi:10.1089/vbz.2011.0631.

2. McIntosh B. Usutu (sa ar 1776), nouvel arbovirus du groupe b. Int Cat Arboviruses 1985; 3:1059–1060.

3. Weissenböck H, Kolodziejek J, Url A, Lussy H, Rebel-Bauder B, Nowotny N. Emergence of Usutu virus, an African mosquito-borne flavivirus of the Japanese encephalitis virus group, central Europe. Emerg Infect Dis. 2002;8(7):652–6. doi:10.3201/eid0807.020094.

4. Buckley A, Dawson A, Gould EA. Detection of seroconversion to West Nile virus, Usutu virus and Sindbis virus in UK sentinel chickens. Virol J. 2006;3:71. doi:10.1186/1743-422X-3-71.5.

5. Rizzoli A, Rosà R, Rosso F, Buckley A, Gould E. West Nile virus circulation detected in northern Italy in sentinel chickens. Vector Borne Zoonotic Dis. 2007;7(3):411–7. doi:10.1089/vbz.2006.0626.

6. Bakonyi T, Erdélyi K, Ursu K, Ferenczi E, Csörgo T, Lussy H, et al. Emergence of Usutu virus in Hungary. J Clin Microbiol. 2007;45(12):3870–4. doi:10.1128/JCM.01390-07.

7. Hubálek Z, Wegner E, Halouzka J, Tryjanowski P, Jerzak L, Sikutová S, et al. Serologic survey of potential vertebrate hosts for West Nile virus in Poland. Viral Immunol. 2008;21(2):247–53. doi:10.1089/vim.2007.0111.

8. Busquets N, Alba A, Allepuz A, Aranda C, Ignacio Nuñez J. Usutu virus sequences in Culex pipiens (Diptera: Culicidae), Spain. Emerg Infect Dis. 2008;14(5):861–3. doi:10.3201/eid1405.071577.

9. Jöst H, Bialonski A, Maus D, Sambri V, Eiden M, Groschup MH, et al. Isolation of usutu virus in Germany. Am J Trop Med Hyg. 2011;85(3):551–3. doi:10.4269/ajtmh.2011.11-0248

10. Vilibic-Cavlek T, Kaic B, Barbic L, Pem-Novosel I, Slavic-Vrzic V, Lesnikar V, et al. First evidence of simultaneous occurrence of West Nile virus and Usutu virus neuroinvasive disease in humans in Croatia during the 2013 outbreak. Infection. 2014;42(4):689–95. doi:10.1007/s15010-014-0625-1.

11. Becker N, Jöst H, Ziegler U, Eiden M, Höper D, Emmerich P, et al. Epizootic emergence of Usutu virus in wild and captive birds in Germany. PLoS One. 2012;7(2):e32604. doi:10.1371/journal.pone.0032604.

12. Gaibani P, Pierro A, Alicino R, Rossini G, Cavrini F, Landini MP, et al. Detection of Usutu-virus-specific IgG in blood donors from northern Italy. Vector Borne Zoonotic Dis. 2012;12(5):431–3. doi:10.1089/vbz.2011.0813.

13. Lupulovic D, Martín-Acebes MA, Lazic S, Alonso-Padilla J, Blázquez AB, Escribano-Romero E, et al. First serological evidence of West Nile virus activity in horses in Serbia. Vector Borne Zoonotic Dis. 2011;11(9):1303–5. doi:10.1089/vbz.2010.0249.

14. Ben Hassine T, De Massis F, Calistri P, Savini G, BelHaj Mohamed B, Ranen A, et al. First detection of co-circulation of West Nile and Usutu viruses in equids in the south-west of Tunisia. Transbound Emerg Dis. 2014;61(5):385–9. doi:10.1111/tbed.12259.

15. Cavrini F, Gaibani P, Longo G, Pierro AM, Rossini G, Bonilauri P, et al. Usutu virus infection in a patient who underwent orthotropic liver transplantation, Italy, August-September 2009. Euro Surveill. 2009;14(50).

16. Pecorari M, Longo G, Gennari W, Grottola A, Sabbatini A, Tagliazucchi S, et al. First human case of Usutu virus neuroinvasive infection, Italy, August-September 2009. Euro Surveill. 2009;14(50).

17. Gaibani P, Pierro AM, Cavrini F, Rossini G, Landini MP, Sambri V. False-positive transcription-mediated amplification assay detection of West Nile virus in blood from a patient with viremia caused by an Usutu virus infection. J Clin Microbiol. 2010;48(9):3338–9. doi:10.1128/JCM.02501-09.

18. Lanciotti RS, Kosoy OL, Laven JJ, Velez JO, Lambert AJ, Johnson AJ, et al. Genetic and serologic properties of Zika virus associated with an epidemic, Yap State, Micronesia, 2007. Emerg Infect Dis. 2008;14(8):1232–9. doi:10.3201/eid1408.080287.

19. Haddow AD, Schuh AJ, Yasuda CY, Kasper MR, Heang V, Huy R, et al. Genetic characterization of Zika virus strains: geographic expansion of the Asian lineage. PLoS Negl Trop Dis. 2012;6(2):e1477. doi:10.1371/journal.pntd.0001477.

20. Duffy MR, Chen TH, Hancock WT, Powers AM, Kool JL, Lanciotti RS, et al. Zika virus outbreak on Yap Island, Federated States of Micronesia. N Engl J Med. 2009;360(24):2536–43. doi:10.1056/NEJMoa0805715.

21. Zanluca C, de Melo VC, Mosimann AL, Dos Santos GI, Dos Santos CN, Luz K. First report of autochthonous transmission of Zika virus in Brazil. Mem Inst Oswaldo Cruz. 2015;110(4):569–72. doi:10.1590/0074-02760150192.

22. Fagbami AH. Zika virus infections in Nigeria: virological and seroepidemiological investigations in Oyo State. J Hyg (Lond). 1979;83(2):213–9.

23. Marchette NJ, Garcia R, Rudnick A. Isolation of Zika virus from Aedes aegypti mosquitoes in Malaysia. Am J Trop Med Hyg. 1969;18(3):411–5. 21.

24. Akoua-Koffi C, Diarrassouba S, Bénié VB, Ngbichi JM, Bozoua T, Bosson A, et al. [Investigation surrounding a fatal case of yellow fever in Côte d'Ivoire in 1999]. Bull Soc Pathol Exot. 2001;94(3):227–30.

25. McCrae AW, Kirya BG. Yellow fever and Zika virus epizootics and enzootics in Uganda. Trans R Soc Trop Med Hyg. 1982;76(4):552–62.

26. Hayes EB. Zika virus outside Africa. Emerg Infect Dis. 2009;15(9):1347–50. doi:10.3201/eid1509.090442.

27. Henderson BE, Hewitt LE, Lule M. Serology of wild mammals. Virus Research Institute Annual Report. East African Printer, Nairobi, Kenya. 1968; 48–51.

28. Darwish MA, Hoogstraal H, Roberts TJ, Ahmed IP, Omar F. A sero-epidemiological survey for certain arboviruses (Togaviridae) in Pakistan. Trans R Soc Trop Med Hyg. 1983;77(4):442–5

29. Heang V, Yasuda CY, Sovann L, Haddow AD, Travassos da Rosa AP, Tesh RB, et al. Zika virus infection, Cambodia, 2010. Emerg Infect Dis. 2012;18(2):349–51. doi:10.3201/eid1802.111224.

30. Foy BD, Kobylinski KC, Chilson Foy JL, Blitvich BJ, Travassos da Rosa A, Haddow AD, et al. Probable non-vector-borne transmission of Zika virus, Colorado, USA. Emerg Infect Dis. 2011;17(5):880–2. doi:10.3201/eid1705.101939.

31. Buckley A, Gould EA. Detection of virus-specific antigen in the nuclei or nucleoli of cells infected with Zika or Langat virus. J Gen Virol. 1988;69 (Pt 8):1913–20. doi:10.1099/0022-1317-69-8-1913.

32. Pan American Health Organization (PAHO). Neurological syndrome, congenital malformations, and Zika virus infection. Implications for public health in the Americas— epidemiological alert. 1 Dec 2015. Available: www.paho.org/hq/index.php?option=com_topics&view=article&id=427&Itemid=41484&lang=en.

33. Oliveira Melo AS, Malinger G, Ximenes R, Szejnfeld PO, Alves Sampaio S, Bispo de Filippis AM. Zika virus intrauterine infection causes fetal brain abnormality and microcephaly: tip of the iceberg? Ultrasound Obstet Gynecol. 2016;47(1):6–7. doi:10.1002/uog.15831.

34. European Centre for Disease Prevention and Control. Rapid risk assessment: Zika virus epidemic in the Americas: potential association with microcephaly and Guillain-Barré syndrome – 10 December 2015. Stockholm: ECDC; 2015.Available: http://ecdc.europa.eu/en/publications/Publications/zika-virus-americas-association-with-microcephaly-rapid-risk-assessment.pdf

35. Kuno G, Chang GJ. Full-length sequencing and genomic characterization of Bagaza, Kedougou, and Zika viruses. Arch Virol. 2007;152(4):687–96. doi:10.1007/s00705-006-0903-z.

36. Pătraşcu IV. Bovine leukemia virus. VII. In vitro replication of virus in bat lung cell culture NBL BLV 2. Virologie. 1988;39(3):199–205.

37. Foster DN, Foster LK. Immortalized cell lines for virus growth. U.S. Pat. No. 5,672,485. 1997.

38. Pasternak AS, Miller WM. First-order toxicity assays for eye irritation using cell lines: parameters that affect in vitro evaluation. Fundam Appl Toxicol. 1995;25(2):253–63.

39. Edgell CJ, McDonald CC, Graham JB. Permanent cell line expressing human factor VIII-related antigen established by hybridization. Proc Natl Acad Sci U S A. 1983;80(12):3734–7.

40. Crandell, R.; Fabricant, C.; Nelson-Rees, W., Development, characterization, and viral susceptibility of a feline (felis catus) renal cell line (crfk). In Vitro 1973, 9, 176–185.

41. Rhim JS, Ro HS, Kim EB, Gilden RV, Huebner RJ. Transformation of horse skin cells by type-C sarcoma viruses. Int J Cancer. 1975;15(2):171–9.

42. Matsuda M, Matsuda N, Watanabe A, Fujisawa R, Yamamoto K, Masuda M. Cell cycle arrest induction by an adenoviral vector expressing HIV-1 Vpr in bovine and feline cells. Biochem Biophys Res Commun. 2003;311(3):748–53.

43. Löffler S, Lottspeich F, Lanza F, Azorsa DO, ter Meulen V, Schneider-Schaulies J. CD9, a tetraspan transmembrane protein, renders cells susceptible to canine distemper virus. J Virol. 1997;71(1):42–9.

44. Shukla P, Nguyen HT, Torian U, Engle RE, Faulk K, Dalton HR, et al. Cross-species infections of cultured cells by hepatitis E virus and discovery of an infectious virus-host recombinant. Proc Natl Acad Sci U S A. 2011;108(6):2438–43. doi:10.1073/pnas.1018878108.

45. Koyama H, Goodpasture C, Miller MM, Teplitz RL, Riggs AD. Establishment and characterization of a cell line from the American opossum (Didelphys virginiana). In Vitro. 1978;14(3):239–46.

46. Amborski R, LoPiccolo G, Amborski G. Development of an established cell line derived from dasypus novemcinctus (armadillo), a laboratory animal susceptible to infection by mycobacterium leprae. Experientia 1974;30: 546–548.

47. Pirtle EC, Woods LK. Cytogenetic alterations in swine kidney cells persistently infected with hog cholera virus and propagated with and without antiserum in the medium. Am J Vet Res. 1968;29(1):153–64.

48. Hull RN, Cherry WR, Tritch OJ. Growth characteristics of monkey kidney cell strains LLC-MK1, LLC-MK2, and LLC-MK2(NCTC-3196) and their utility in virus research. J Exp Med. 1962;115:903–18.

49. McClurkin AW, Pirtle EC, Coria MF, Smith RL. Comparison of low-and high-passage bovine turbinate cells for assay of bovine viral diarrhea virus. Arch Gesamte Virusforsch. 1974;45(3):285–9.

49. Gaush CR, Hard WL, Smith TF. Characterization of an established line of canine kidney cells (MDCK). Proc Soc Exp Biol Med. 1966;122(3):931–5.

50. Schechter EM, Summers J, Ogston CW. Characterization of a herpesvirus isolated from woodchuck hepatocytes. J Gen Virol. 1988;69 (Pt 7):1591–9. doi:10.1099/0022-1317-69-7-1591.

51. Henderson IC, Lieber MM, Todaro GJ. Mink cell line Mv 1 Lu (CCL 64). Focus formation and the generation of "nonproducer" transformed cell lines with murine and feline sarcoma viruses. Virology. 1974;60(1):282–7.

52. Acosta EG, Castilla V, Damonte EB. Alternative infectious entry pathways for dengue virus serotypes into mammalian cells. Cell Microbiol. 2009;11(10):1533–49. doi:10.1111/j.1462-5822.2009.01345.x

53. Thaisomboonsuk BK, Clayson ET, Pantuwatana S, Vaughn DW, Endy TP. Characterization of dengue-2 virus binding to surfaces of mammalian and insect cells. Am J Trop Med Hyg. 2005;72(4):375–83.

54. Liao M, Kielian M. Domain III from class II fusion proteins functions as a dominant-negative inhibitor of virus membrane fusion. J Cell Biol. 2005;171(1):111–20. doi:10.1083/jcb.200507075.

55. Bakonyi T, Lussy H, Weissenböck H, Hornyák A, Nowotny N. In vitro host-cell susceptibility to Usutu virus. Emerg Infect Dis. 2005;11(2):298–301. doi:10.3201/eid1102.041016.

56. Barr KL, Anderson BD, Heil GL, Friary JA, Gray GC, Focks DA. Dengue serotypes 1–4 exhibit unique host specificity in vitro. Virus Adaptation and Treatment. 2012; 4:65–73. doi: http://dx.doi.org/10.2147/VAAT.S36856

57. Barr KL, Anderson BD. Dengue viruses exhibit strain-specific infectivity and entry requirements in vitro. Virus Adaptation and Treatment. 2013; 5:1–9. doi: http://dx.doi.org/10.2147/VAAT.S38143

58. Burke D, Monath T. Flaviviruses. In: Knipe D, Howley P, Editors. Fields Virology 4 ed. Philadelphia: Lippincott Williams & Wilkins; 2001. Vol. 1, pp 1043–1126.

59. van der Schaar HM, Rust MJ, Chen C, van der Ende-Metselaar H, Wilschut J, Zhuang X, et al. Dissecting the cell entry pathway of dengue virus by single-particle tracking in living cells. PLoS Pathog. 2008;4(12):e1000244. doi:10.1371/journal.ppat.1000244.

60. Ishak R, Tovey DG, Howard CR. Morphogenesis of yellow fever virus 17D in infected cell cultures. J Gen Virol. 1988;69 (Pt 2):325–35. doi:10.1099/0022-1317-69-2-325.

61. Ng ML, Lau LC. Possible involvement of receptors in the entry of Kunjin virus into Vero cells. Arch Virol. 1988;100(3-4):199–211.

62. Nawa M, Takasaki T, Yamada K, Kurane I, Akatsuka T. Interference in Japanese encephalitis virus infection of Vero cells by a cationic amphiphilic drug, chlorpromazine. J Gen Virol. 2003;84(Pt 7):1737–41. doi:10.1099/vir.0.18883-0.

63. Harrison SC. The pH sensor for flavivirus membrane fusion. J Cell Biol2008;183(2):177–9. doi:10.1083/jcb.200809175.

64. Fritz R, Stiasny K, Heinz FX. Identification of specific histidines as pH sensors in flavivirus membrane fusion. J Cell Biol. 2008;183(2):353–61. doi:10.1083/jcb.200806081.

65. Stiasny K, Fritz R, Pangerl K, Heinz FX. Molecular mechanisms of flavivirus membrane fusion. Amino Acids. 2011;41(5):1159–63. doi:10.1007/s00726-009-0370-4.

66. Mukhopadhyay S, Kuhn RJ, Rossmann MG. A structural perspective of the flavivirus life cycle. Nat Rev Microbiol. 2005;3(1):13–22. doi:10.1038/nrmicro1067.

67. Smit JM, Moesker B, Rodenhuis-Zybert I, Wilschut J. Flavivirus cell entry and membrane fusion. Viruses. 2011;3(2):160–71. doi:10.3390/v3020160.

68. Holzmann H, Heinz FX, Mandl CW, Guirakhoo F, Kunz C. A single amino acid substitution in envelope protein E of tick-borne encephalitis virus leads to attenuation in the mouse model. J Virol. 1990;64(10):5156–9.

69. Lobigs M, Usha R, Nestorowicz A, Marshall ID, Weir RC, Dalgarno L. Host cell selection of Murray Valley encephalitis virus variants altered at an RGD sequence in the envelope protein and in mouse virulence. Virology. 1990;176(2):587–95.

70. Cecilia D, Gould EA. Nucleotide changes responsible for loss of neuroinvasiveness in Japanese encephalitis virus neutralization-resistant mutants. Virology. 1991;181(1):70–7.

71. Gritsun TS, Lashkevich VA, Gould EA. Nucleotide and deduced amino acid sequence of the envelope glycoprotein of Omsk haemorrhagic fever virus; comparison with other flaviviruses. J Gen Virol. 1993;74 (Pt 2):287–91. doi:10.1099/0022-1317-74-2-287.

72. Bakonyi T, Gould EA, Kolodziejek J, Weissenböck H, Nowotny N. Complete genome analysis and molecular characterization of Usutu virus that emerged in Austria in 2001: comparison with the South African strain SAAR-1776 and other flaviviruses. Virology. 2004;328(2):301–10. doi:10.1016/j.virol.2004.08.005.

73. Peletto S, Lo Presti A, Modesto P, Cella E, Acutis PL, Farchi F, et al. Genetic diversity of Usutu virus. New Microbiol. 2012;35(2):167–74.

74. Chambers TJ, Halevy M, Nestorowicz A, Rice CM, Lustig S. West Nile virus envelope proteins: nucleotide sequence analysis of strains differing in mouse neuroinvasiveness. J Gen Virol. 1998;79 ( Pt 10):2375–80. doi:10.1099/0022-1317-79-10-2375.

75. Kirkham M, Parton RG. Clathrin-independent endocytosis: new insights into caveolae and non-caveolar lipid raft carriers. Biochim Biophys Acta. 2005;1746(3):349–63.

76. Marsh M, Helenius A. Virus entry: open sesame. Cell. 2006;124(4):729–40. doi:10.1016/j.cell.2006.02.007.

77. Conner SD, Schmid SL. Regulated portals of entry into the cell. Nature. 2003;422(6927):37–44. doi:10.1038/nature01451.

78. Swanson J, Watts C. Macropinitosis. Trends Cell Biol. 1995; 5: 424–428.

